# Dynamic changes in Anterior Cingulate Cortex ensembles mark the transition from exploration to exploitation

**DOI:** 10.1101/605592

**Authors:** Eldon Emberly, K Seamans Jeremy

## Abstract

The ability to acquire knowledge about the value of stimuli or actions factors into simple foraging behaviors as well as complex forms of decision making. The anterior cingulate cortex (ACC) is thought to be involved in these processes, although the manner in which neural representations acquire value is unclear. Here we recorded from ensembles of ACC neurons as rats learned which of 3 levers was rewarded each day through a trial and error process. Action representations remained largely stable during exploration, but there was an abrupt, coordinated and differential change in the representation of rewarded and nonrewarded levers by ACC neurons at the point where the rat realized which lever was rewarded and began to exploit it. Thus, rather than a gradual, incremental process, value learning in ACC can occur in an all-or-none manner and help to initiate strategic shifts in forging behavior.

## Introduction

Foraging is a central component of the daily life of many species and involves striking a balance between exploiting a known resource and exploring a new one. The transition from exploration to exploitation is based on an evaluation of what options are currently available and most valuable to the organism. While the ‘explore/exploit tradeoff’ is an interesting issue from the point of view of behavioral ecology, the basic processes underlie other more complex forms of behavior and decision making (Addicott et al 2017; Pearson et al 2014).

Foraging decisions involve multiple brain regions including the striatum, midbrain and brain stem neuromodulatory centers and the medial frontal cortex (Aston-Jones & Cohen 2005; Daw et al 2006; Cohen et al 2007; Rushworth & Behrens 2008; Hayden et al 2011, Kolling et al 2012; Mobbs et al 2013; Khani & Rainer 2016). The results of human functional MRI studies suggest that the anterior cingulate cortex (ACC) plays a central role in foraging decisions by virtue of its involvement in the flexible representation of value. In humans, the ACC is activated by the most highly valued option or a comparison of the relative value of available options (Kolling et al 2012; 2016; Rushworth et al, 2012; Basten et al 2010; Klein-Flugge et al 2016; de Berker et al 2019). The ACC may also encode the value of search which in can turn impact decisions about whether to stick with one option or continue to explore (Kolling et al 2012; 2016). A similar picture has emerged from electrophysiology studies in animals as single ACC neurons can represent the relative value of an action or outcome (Pratt & Mizumori 2001; Hayden & Platt 2010; Hillman & Bilkey 2010; Kennerley et al 2009; Wallis & Kennerley 2011; Cai & Padoa-Schioppa 2012; Cowen et al 2012; Narayanan et al 2013; Azab & Hayden 2017; 2018; Hyman et al 2017; Monosov 2017) as well as the relative value of leaving a depleting patch for a new one (Hayden et al, 2011; Blanchard & Hayden 2014).

ACC neurons and ensembles also encode a regency-weighted history of past outcomes (Seo & Lee 2007; Bernacchia et al 2011; Hyman et al 2017) and it has been argued that these signals might produce the value representations that ultimately guide foraging decisions (Kolling et al 2016; Wittmann et al 2016). The idea that action representations accrue value through an experience-dependent process is consistent with theories of reinforcement learning (RL) in mesolimbic pathways where value representations are thought to be continuously updated through a dopamine-dependent process (Stauffer et al 2014; Schultz 2015; Kolling & Akam 2017; Sharpe & Schoenbaum 2018).

In order to understand the neural basis of foraging, it is also important to consider how foraging decisions are made in naturalistic situations. Rats are ‘central-place’ foragers in that once they discover a food cache, they immediately return it to their burrow (Barnett 2005). Since they may not know, or fail to remember which actions led to a cache or where a cache is located, value can only be imbued on an action or location once a cache is discovered. All-or-none value attributions also occur during creative exploration (Bowden et al 2005) in humans. For instance, Hart et al (2017) observed that subjects meandered through a virtual foraging landscape while occasionally making sudden ‘creative leaps’ about ambiguous objects, believing that they were valuable and abruptly redirecting their responses towards them. Such foraging decisions place a high premium on novelty, which by definition cannot be acquired through a gradual process. Therefore, at least in certain situations, an all-or-none value attribution process may drive the shift from exploration to exploitation.

The shift from exploration to exploitation produces an immediate change in perspectives and goals. Within the ACC we have observed that global shifts in ensemble dynamics occur the moment a rat detects that the environment, task or motivational context has changed (Durstewitz et al 2010; Hyman et al 2012; Ma et al 2016; Caracheo et al 2013;2018). These ensemble shifts are associated with an immediate reconfiguration in the way common stimuli and actions are represented (Ma et al 2016). On the other hand, representations remain stable if the overall task or motivational context remains the same (Powell & Redish 2013; 2017; Caracheo et al 2018). Based on these considerations, we would predict that ACC ensembles should track outcomes in a consistent manner during exploratory foraging but undergo a sudden reconfiguration during the shift to exploitation.

To investigate this issue, we recorded ACC neuronal dynamics during a simple operant foraging task where rats sampled from 3 levers to find the one that was rewarded each day. Even though the task was deterministic, there was a large variability in foraging behavior, as some rats discovered and exploited the rewarded lever immediately while others displayed initial lever biases or chose levers randomly throughout the entire session. The present study focused on the subset of sessions where rats exhibited both extensive periods of exploration and exploitation. Arrays of driveable tetrodes recorded multiple single neuron activity in the ACC and the activity surrounding presses on each of the 3 levers was analyzed throughout the task, including the point when the rats transitioned from exploration to exploitation. Most neurons were at least partially responsive to some set of lever presses and the main shift in lever press responses occurred just before the transition from exploration to exploitation.

## Results

### Behavior

The database included 14 free choice sessions from 9 rats. In order to quantify choice behavior, a press on the rewarded lever was scored as a 1 while presses on the 2 non-rewarded levers were scored as −1 and the normalized cumulative sums were plotted (Fig 1). In spite of the fact that the correct lever was rewarded 100% of the time, rats took an average of 70 trials to fully exploit it, owing in part to lever biases from previous sessions. Since a main goal of the study was to understand the neural dynamics associated with the transition from exploration to exploitation, it was important that both strategies appeared in a given session. In sessions 1,2,7 the rats seized on the correct lever early and exhibited very little exploration, while in sessions 4 and 11, the rat’s choices remained essentially random and there was no clear transition to an exploit strategy (gray curves). During the remaining sessions (black curves), the rats went through a variable length exploration period before exploiting the correct lever. As a result, subsequent analyses will focus mainly on these 9 sessions. For each session, a curve was fit to the cumulative sum choice vector and the minima of this curve was taken as the *behavioral transition point,* which demarcated the theoretical transition between explore and exploit foraging strategies. The trials before and after the behavioral transition point will henceforth be referred to as blocks 1 and 2 respectively.

**Figure 1:**
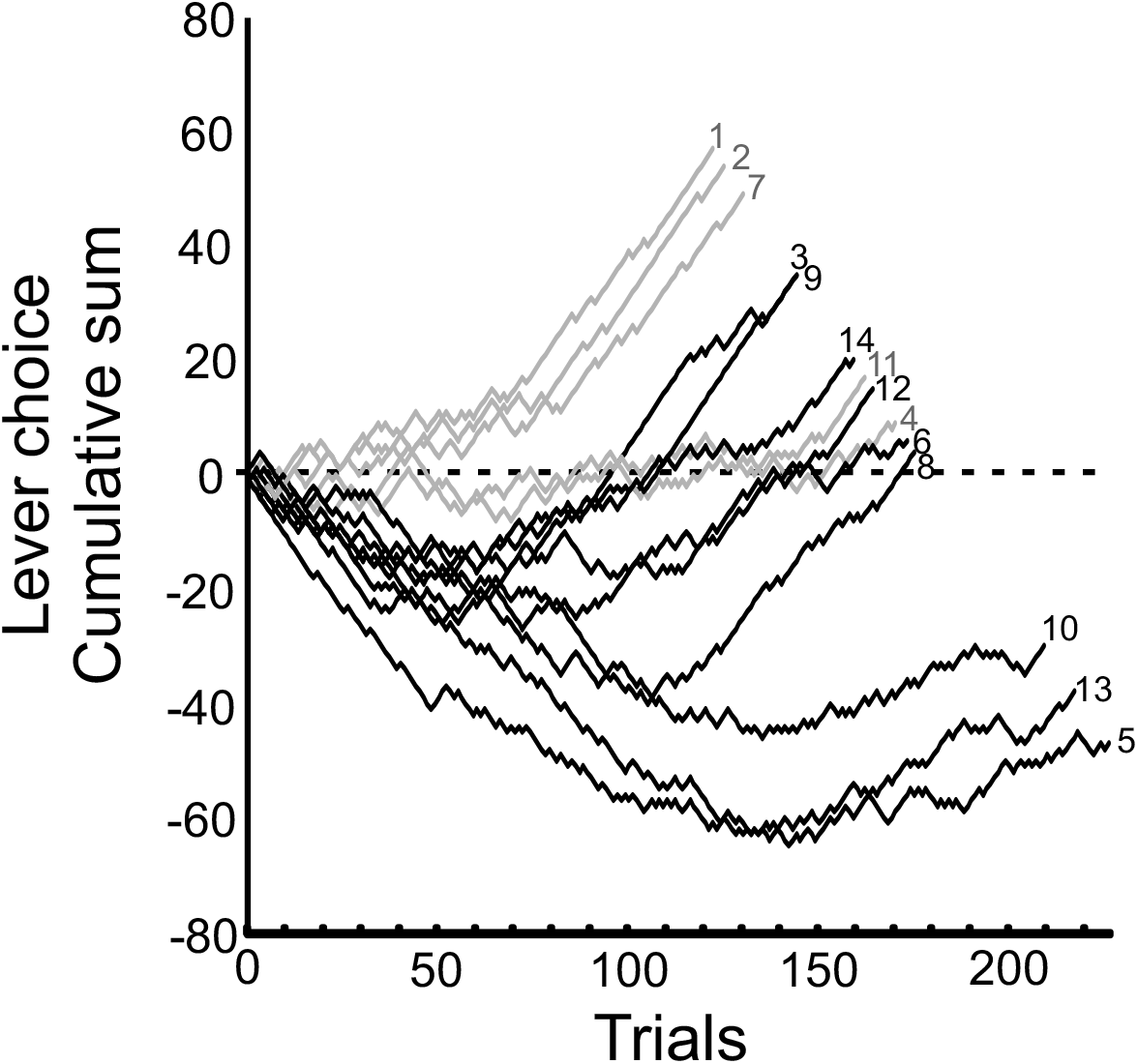
Cumulative distribution of lever presses across sessions. For each session, presses on the rewarded were scored as 1 and presses on rewarded levers as −1 and the cumulative sum was calculated. In sessions 1,2,7 there was little to no exploration phase as rats immediately seized upon the correct lever early in the sessions while in sessions 4 and 11, the rats randomly chose levers throughout and did not exhibit a clear exploitation phase (gray curves). The remaining sessions 3,5,6,8,9,10,12,13,14 (black curves) were characterized by protracted exploration and exploitation phases.

### Neural Dynamics

Most neurons responded to presses on more than one lever. As a result we refrained from categorizing individual neurons as having a lever preference. Instead, we compared how the relative preferences of all neurons for the 3 levers changed in blocks 1 and 2. A t-statistic (t-stat) compared the mean spike counts for each neuron during presses on one lever versus presses on the other 2 levers. A positive t-stat meant that the neuron fired more on average to a given lever than the other 2 levers whereas the opposite was true of a negative t-stat. Since the t-stats were calculated separately for each lever in blocks 1 and 2, it yielded a total of 6 t-stat values/neuron and 2970 in total for the population. The respective positive and negative t-stat values tails in the 95^th^ and 5^th^ percentiles of this distribution were determined. A binomial test assessed whether the number of neurons attaining a t-stat value beyond these thresholds were significant for each lever (assuming that suprathreshold values should occur 5% of the time). Figure 2A plots the number of neurons that attained a suprathreshold t-stat value for each of the 3 levers in blocks 1 and 2. The number of neurons attaining a significant positive or negative t-stat were balanced across the 3 levers in block 1. However, the t-stat distribution shifted after the behavioral transition point as significantly more neurons exhibited significant positive t-stat values for the rewarded lever and significant negative t-stat values for the ‘other’ lever in block 2.

**Figure 2:**
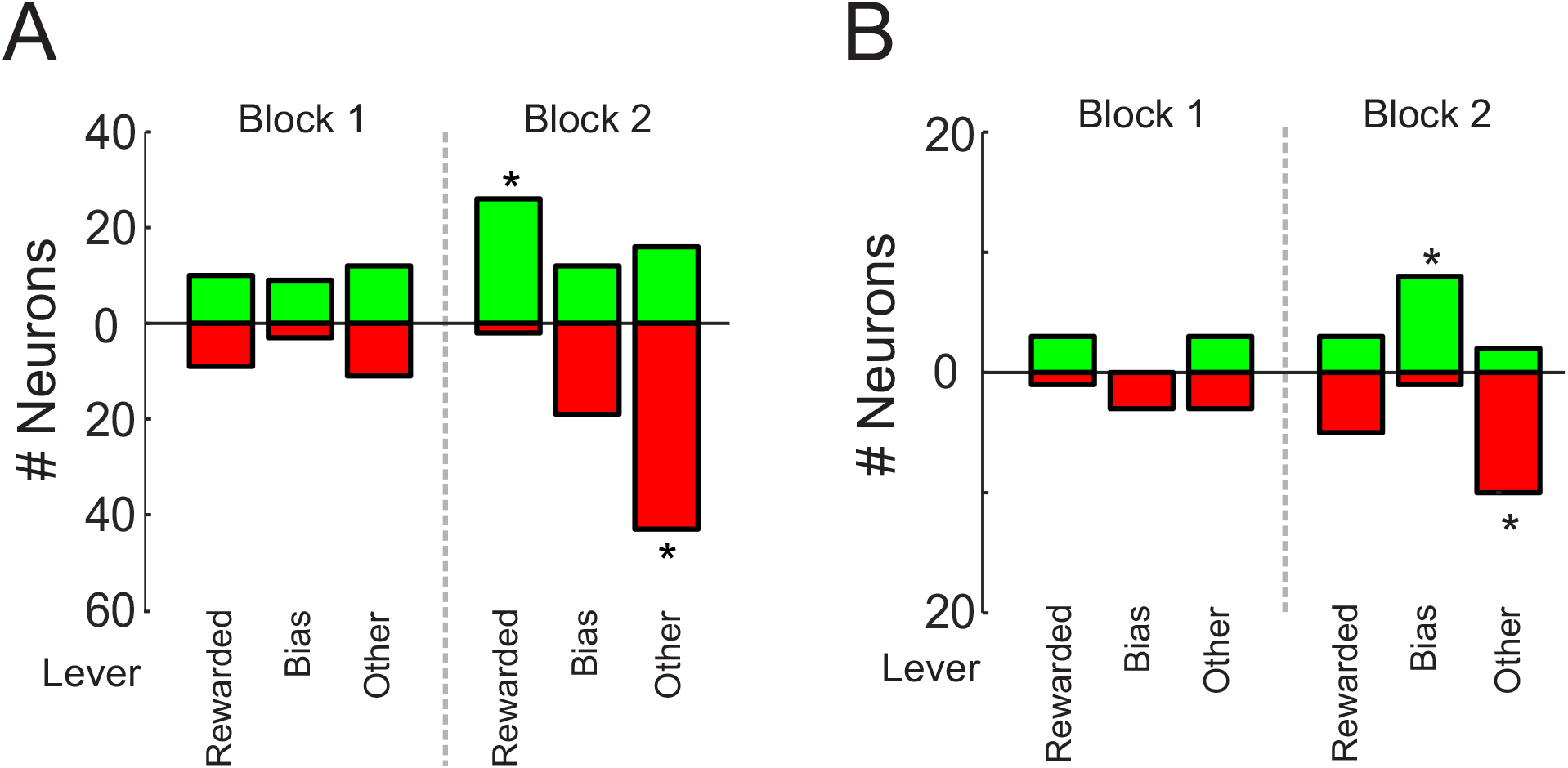
Lever response distributions as assessed using a t-statistic (t-stat). For each neuron, a t-stat compared the mean response on a given lever versus the mean responses on the other 2 levers in blocks 1 and 2. **A)** The distribution of t-stat values for all neurons in sessions 3,5,6,8,9,10,12,13,14 that exhibited a t-stat value in the top 5% of the overall distribution. **B)** The distribution of t-stat values for all neurons in sessions 4 and 11 that exhibited a t-stat value in the top 5% of the overall distribution. Green and red bars denote positive and negative t-tstat values respectively. * denotes that a significantly higher proportion of neurons than expected, exhibited large t-stat values for a given lever. Lever designations were as follows: The ‘rewarded’ lever paid out a single food pellet when pressed, the ‘bias’ lever was the nonrewarded lever a given rat pressed the most in block 1 and the remaining nonrewarded lever was referred to as the ‘other’ lever.

For comparison, the same analysis was performed on data from sessions 4 and 11 where the transition to a successful exploit strategy never occurred (see Fig. 1). The overall pattern was quite similar for these sessions (Fig 2B), including the fact that more neurons exhibited a significant negative t-stat value for the ‘other’ lever in block 2. The main difference was that the number of neurons that exhibited a positive t-stat value for the rewarded lever in block 2 did not show a significant increase.

These analyses suggested that for the 9 sessions of interest, a shift had occurred in the way the neurons responded to the 3 levers after the behavioral transition point. Because the t-stat compared mean spike counts across large numbers of trials, it could not provide information about exactly when the shift occurred. In order to address this question, we separately plotted the normalized, interpolated spike counts of all neurons to rewarded (Fig 3A) and nonrewarded (Fig 3B,C) lever presses throughout the sessions. The neurons were sorted based on the differences in the spike counts in blocks 1 versus 2 and at the top of the distributions were neurons that fired more to a given lever in block 1 than 2 whereas the opposite was true of neurons at the bottom of the distributions. The most obvious changes in lever press responses occurred in close proximity to the behavioral transition point. In Fig 3C the neurons were resorted in the same order as in Fig 3A and while the differential responses at the top and the bottom of the distribution were still apparent, they were not as obvious as in Fig 3B, suggesting that some neurons changed their responses across all levers concurrently while others changed their responses more for one lever than another.

**Figure 3:**
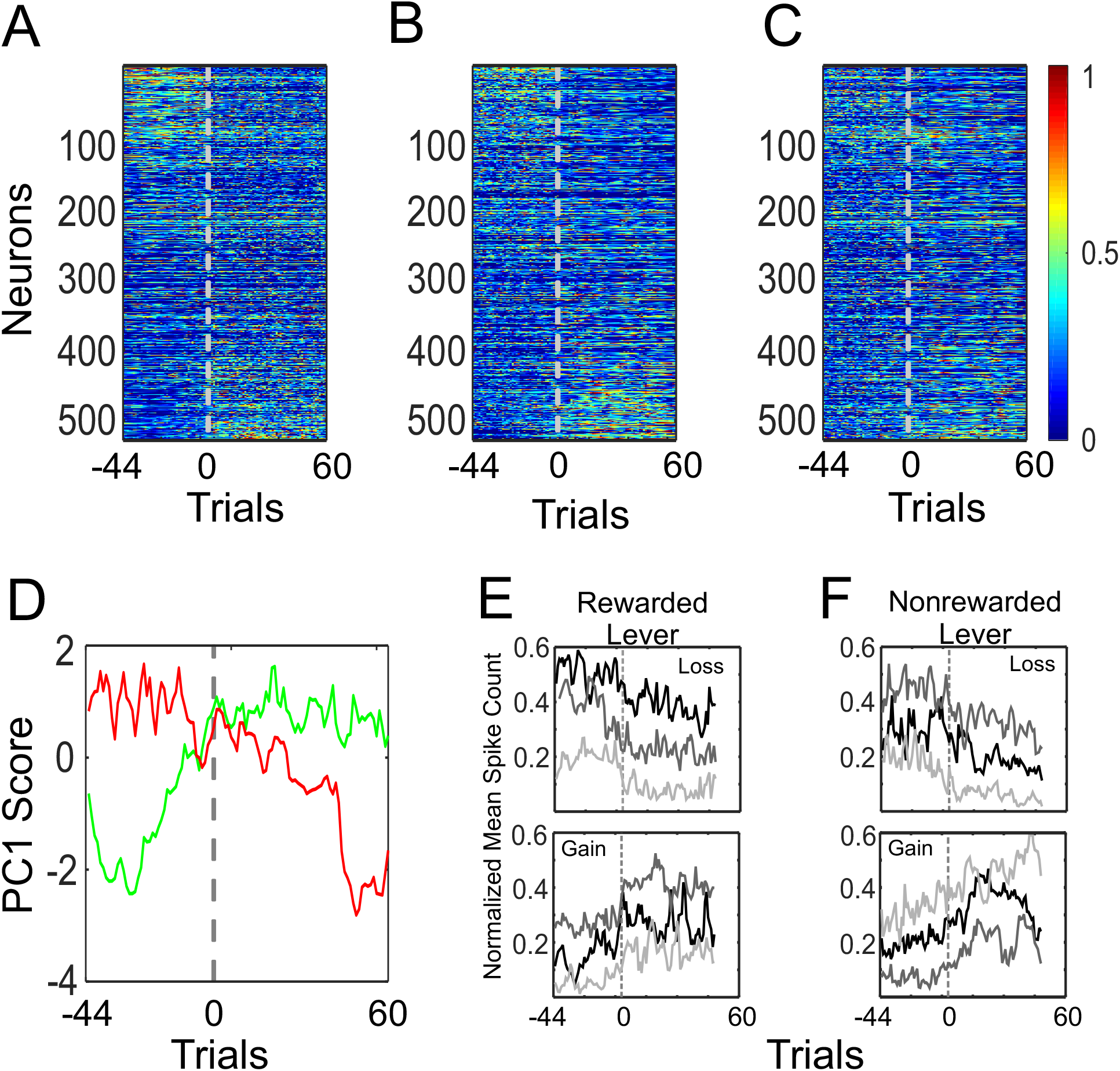
Changes in spike counts at the behavioral change point. The normalized, interpolated spike counts during the 1s period prior to rewarded **(A)** or nonrewarded **(B)** lever presses for all neurons in sessions 3,5,6,8,9,10,12,13,14. Neurons were sorted based on the difference in the mean normalized, interpolated spike counts in block 1 versus block 2. **C**) The normalized, interpolated spike count vectors in B were resorted to match the ordering of neurons in A. **D**) The first principle component (PC1) of the matrix shown in A (green) and the matrix shown in B (red). **E**) The 100 neurons in the top and bottom of the matrix shown in A were clustered using k-means. The centers of the 3 resultant clusters for the top and bottom 100 neurons are shown in the top and bottom panels respectively. **F**) Same as E but for the 100 neurons in the top and bottom of the matrix shown in B. The gray dotted line denotes the behavioral transition point to which all the normalized, interpolated spike count vectors were aligned. Because the spike count vectors were normalized, the y-axis gives the proportion of change out of a maximum of 1.

The matrices in Figs 3A and B were then subjected to Principal Component Analysis (PCA). PC1 captured the larges change in rewarded lever responses that peaked at the behavioral transition point (Fig 3D, green curve). PC1 of the nonrewarded lever responses remained steady until just before the behavioural transition point at which point it exhibited a smaller shift followed by a slow drift, culminating in a final late drop (Fig 3D, red curve).

While PCA was a useful way to show overall trends in the population, the main changes of interest occurred mainly in the groups of neurons at the top and bottom of the distributions in Figs 3A,B. Therefore, we extracted the top and bottom 100 neurons and used k-means clustering to better characterize the neural transitions. The mean changes (i.e. the centers of each cluster) in the responses to rewarded and nonrewarded lever presses are shown in Figures 3E and F respectively. In all 4 subgroups of neurons, the losses or gains in lever press responses occurred at, or just before the behavioural transition point. The only exception being the more gradual and delayed changes by neurons that exhibited gains in nonrewarded lever press responses (Fig 3F, bottom).

An even finer perspective on the underlying dynamics is provided by the single neuron examples in Fig 4. The neurons in Figs. 4A and B responded to both rewarded and non-rewarded lever presses in block 1 but only the responses to the rewarded lever increased after the behavioral transition point. The neuron in Fig 4C had a high basal firing rate but showed little modulation at the time of the lever presses in block 1, yet the rewarded lever presses selectively increased in block 2. In Fig 4D, the neuron exhibited almost no lever-press responsivity in block 1 but became strongly lever-press responsive in block 2.

**Figure 4:**
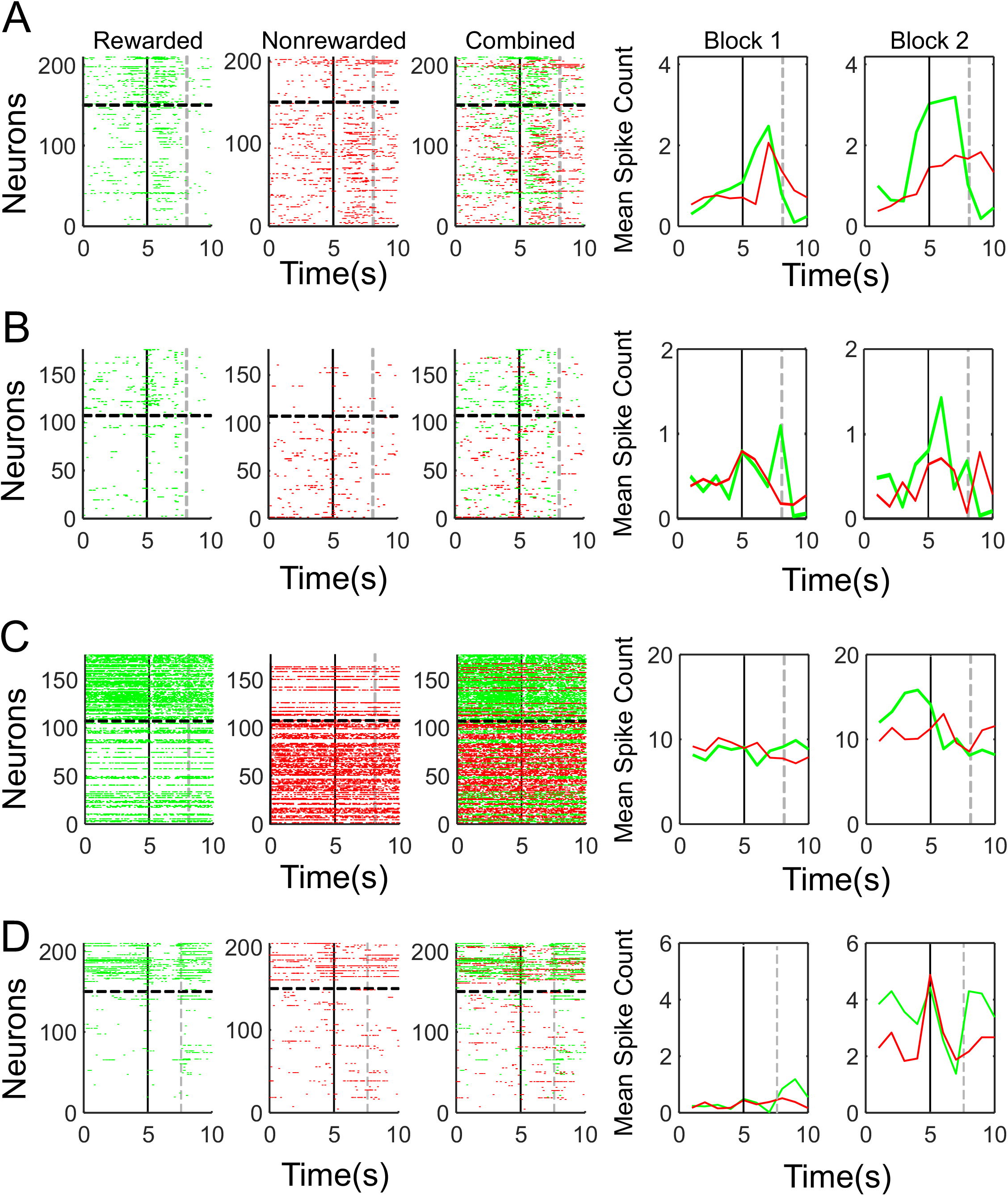
Individual examples of block-dependent changes in lever press responses (**A-D**). In each series, the far left panel is the raster of all spikes during the 10s period centered on rewarded lever presses (green), the middle left panel is the raster of all spikes during the 10s period centered on nonrewarded lever presses (red), while the middle right panel combines these two rasters. The two panels at the far right are the overall mean lever press responses for block 1 (left) and block 2 (right) for rewarded lever presses (green) and nonrewarded lever presses (red). In each panel the black, solid horizontal line separates blocks 1 and 2. The black, solid vertical line is the time of the lever press and the light gray dotted, vertical line is the time that the outcome was delivered on rewarded trial or the time the outcome would have been delivered on nonrewarded trials.

In some neurons, the change in lever press responses was associated with a change in background firing. To visualize this, for each neuron all time bins outside of the trials (i.e. all timebins excluding those 1 s prior to levers-out to 1 s after the outcome) were extracted and concatenated into a continuous vector that was aligned to the behavioral change point. The matrix of these vectors was sorted such that neurons with the largest decrease in spiking from blocks 1 to 2 were at the top and those with the largest increase, at the bottom (Fig. 5A). Although the change in PC1 of this matrix was not as abrupt or synchronized as the changes shown in Fig. 3B, changes in background activity appeared throughout block 1 which were largely absent in block 2 (Fig 5B). Based on the results of two-sample Kolmogorov-Smirnov tests, 130 neurons exhibited a significant difference in baseline spike counts in blocks 1 versus 2. However, across all neurons, a significant correlation existed between the block 1 to block 2 change in spiking during the background and rewarded lever press periods (r=0.36, p<0.0001) and between the background and nonrewarded lever press periods (r=0.34, p<0.0001). Examples of neurons exhibiting concurrent changes during the background and lever-press periods are provided in Fig. 5C-F.

**Figure 5:**
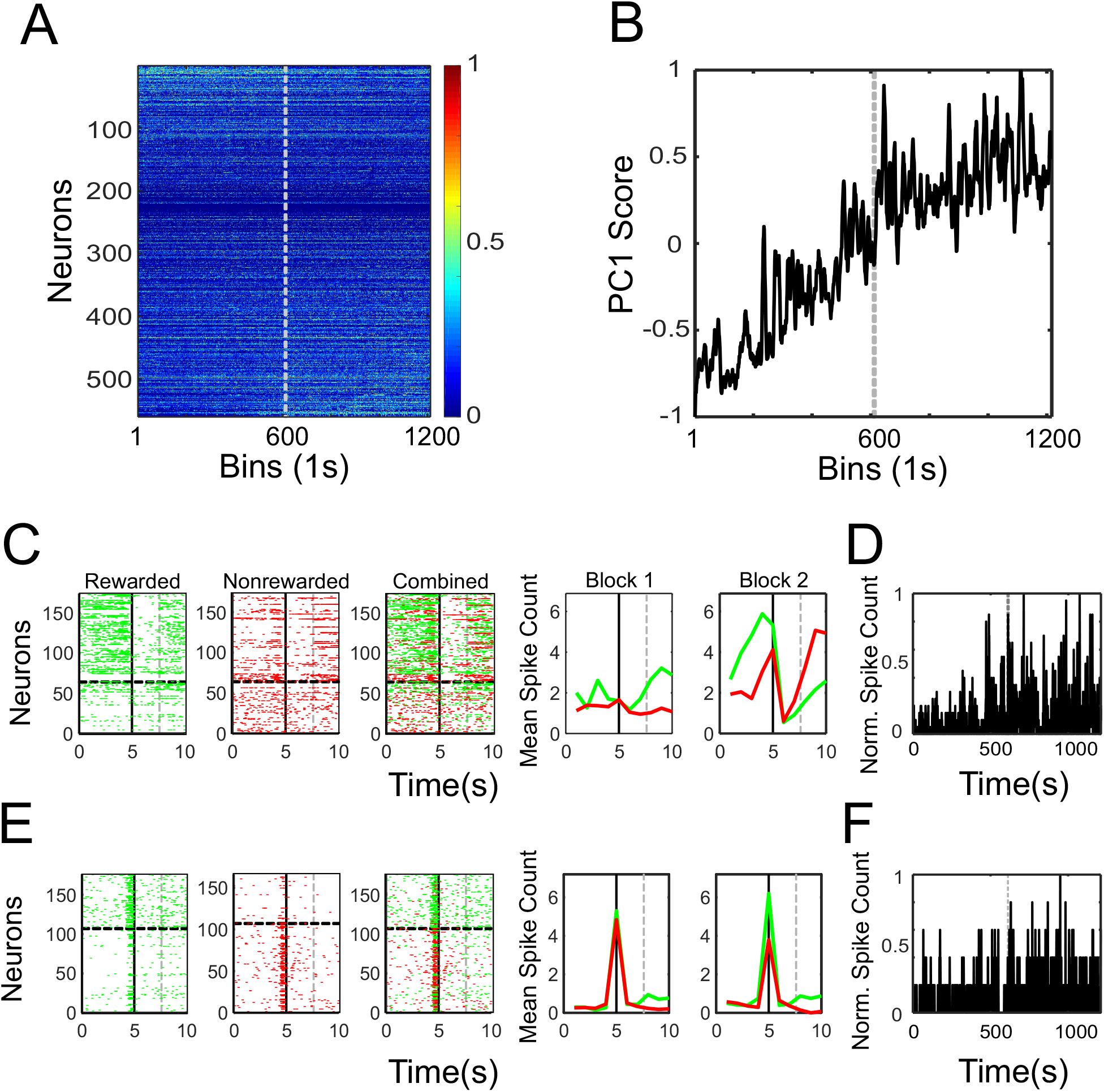
Changes in background activity. **A)** The normalized, interpolated spike counts for all timebins (1s) throughout the session excluding the times surrounding the lever press and outcome periods (i.e. the ‘background period’). The neurons were sorted based on the difference in mean spike counts during the background period of blocks 1 versus 2. The gray dotted line is the behavioral transition point separating blocks 1 and 2. **B)** The first principle component (PC1) of the matrix shown in A. **C)** Example of block-dependent changes in the lever press responses of an individual neuron. The panel layout is identical to Figure 4. **D)** Changes in the normalized spike counts during the background period for the neuron shown in C. The gray vertical line denotes the behavioral change point. **(E-F)** A second example neuron.

## Discussion

In the present study we found that there was a change in lever-press responses across the population of ACC neurons during the switch from exploration to exploitation on a simple 3-lever foraging task. Overall, more neurons began to respond at a relatively higher rate to the rewarded lever and a relatively lower rate to a nonrewarded lever near the behavioral transition point. In addition, approximately one quarter of the population exhibited changes in baseline activity around the behavioral transition point and the changes in activity during baseline and lever press periods were correlated. Collectively the results indicated that the change from exploration to exploitation was marked by a coordinated reconfiguration of ACC activity patterns both inside and outside of the lever-press periods.

Rasters of individual ACC neurons can give the impression that these neurons are inherently tuned to a certain task element, such as a lever press. However, ACC neurons are both highly multi-responsive and highly variable in their responses from trial-to-trial (Hayden & Platt 2010; Ma et al 2016). ACC neurons can exhibit responses to a diverse mixture of task elements which can remain quite stable if the task parameters and the physical environment do not change. On the other hand, neurons that exhibit strong responses to a given task element in one context can be non-responsive to that element and/or responsive to another task element when the physical or task context is changed (Durstewitz et al 2010; Hyman et al 2012; Ma et al 2016; Malagon-Vina et al 2018). These changes tend to be abrupt and are felt to varying degrees across the neuronal population (Hyman et al 2012; Caracheo et al 2013; Ma et al 2016). Nevertheless, they occur in a very well-balanced manner, such that the overall level of activity and even the overall amount of task-responsivity remains essentially constant (Ma et al 2016). In other words, a change in context appears to redistribute a fixed amount of task responsivity across a population of neurons.

These contextual shifts in activity are probably not directly driven by sensory information but reflect underlying changes in the animal’s internal state. The ACC is a central hub in both the Papez and limbic circuits and is extensively interconnected with regions involved in somatovisceral processing (Groenewegen et al 1999; 2000; Van Eden et al 2000). The ACC is therefore anatomically positioned to both receive information about internal state as well as directly influence it (Neafsey 1990; Feldman & Conforti 1985; Critchley et al 2003). We recently observed a particularly striking example of the impact of changes in internal state on ACC ensembles as statistically separable activity-state patterns emerged in a single chamber in response to blocks of aversive, neutral and appetitive events. The ensembles shifted to a given state abruptly but once initiated, the states were maintained long after the initiating events had terminated (Caracheo et al 2013; 2018). The ensemble dynamics associated with switching affective contexts shared many similarities with the shifts that occurred when animals were switched between physical or task contexts. They were also similar to the shifts in ensemble dynamics observed in the present study, probably because the transition from exploration to exploitation is associated with a significant change in the animal’s perspectives, goals and internal state.

Other factors likely also contributed the changes in ensemble dynamics from blocks 1 to 2. Rat medial frontal cortex neurons track changes in body positions or paths traveled (Euston et al 2006; Cowen et al 2007) and therefore, any differences in the way the animals interacted with the levers in blocks 1 and 2 could have potentially impacted the neuronal responses. Yet we have found that within the confines of a small operant chamber, the encoding of limb positions or movements, while present, tend to be weak for most ACC neurons (Lindsay et al 2018). This would be especially true when animals are interacting with an identical lever, in the identical location in two task epochs as was the case in the present study. It is therefore unlikely that the encoding of movement differences could fully account for the large and widespread redistribution of responsivity reported here.

After the behavioral transition point, the animal shifted their responses to the rewarded lever and therefore it could be argued that the behavioral transition point was where the rewarded lever (or lever-press action) had become imbued with value. RL provides a conceptual framework for explaining how value learning may occur in other brain regions (Schultz 2015; Sharpe & Schoenbaum 2018). The main premise of RL is that the goal of the agent (in this case, the rat) is to maximize total rewards (Sutton & Barto 1998). The value function and the action policy that seeks to maximize it are updated in a gradual, iterative manner.

The abruptness of the change in the response to the rewarded and nonrewarded lever presses therefore seems at odds with a gradual RL process. Instead, it is more consistent with learning by insight, where the optimal solution is acquired all at once. Of course the only way the animal could achieve this type of insight would be by keeping a tally of past outcomes. It is worth noting that ACC neurons also track recent outcomes and a neural trace of past outcomes can persist into subsequent trials (Seo & Lee 2007; Bernacchia et al 2011; Hyman et al 2017). It has been suggested that these outcome representations could in theory be used to generate the value estimates that ultimately guide foraging decisions (Kolling et al 2016; Wittmann et al 2016). If we assume that the point where the rats realized a lever had value was the point where they began to exploit it, then we may also conclude that the coincident shifts in lever responses were an integral part of the valuation process. Hunt et al (2018) recently suggested that the ACC integrates various types of information towards a decision bound that once reached, evokes an update in belief structures and associated actions. To put this in a mechanistic framework, we would argue that subsets of ACC neurons integrate information about past outcomes (Seo & Lee 2007; Hyman et al 2017) until a decision bound is reached at which point action or stimulus representations acquire value in an all-or-none manner. The associated shift in ensemble dynamics then initiates changes in internal state that signals that something is fundamentally different and now may be the time to consider updating the current goals or strategies.

## Methods

### Behavior

The choices of the animals were scored as either a 1 or −1 depending or whether a choice was correct or incorrect. A sine function was fit to the cumulative sum of these binary values and the minima of this curve was designated as the behavioral transition point. A total of 9 sessions were chosen that had at least 44 trials before (block 1) and 60 trials after (block 2) the behavioral transition point. These 9 sessions were aligned to the behavioral transition point and trials more than 44 before the behavioral transition point in block 1 or more than 60 after the behavioral transition point in block 2, were excluded. Levers were always labeled as follows (regardless of their physical location): The lever that was rewarded was referred to as the ‘rewarded’ lever, the nonrewarded lever that the rat presses most often in block 1 was referred to as the ‘bias’ lever and the remaining nonrewarded lever was referred to as the ‘other’ lever.

### Neural analyses

The total database included 825 neurons recorded across 14 sessions from 9 rats. This population was reduced to 595 neurons in the 9 sessions of interest. A t-statistic (t-stat) that compared lever press responses was calculated on the 1 sec. time bin immediately preceding each lever press. In order to ensure there was sufficient data to calculate a t-stat, neurons had to have a mean spike count of at least 0.2 spikes/bin across all trials on at least one of the levers. This further restricted the total neurons from 595 to 495 for the 9 sessions of interest. The t-stat was calculated for each lever in each block and compared the mean spike counts associated with all presses on a given lever versus the mean spike counts associated with all presses on the other 2 levers. When computed across all neurons, levers and blocks, this yielded a total of 2970 t-stat values. We then identified the positive and negative t-stat values that resided at the top and bottom 5% of the distribution. Neurons with a t-stat value above or below these thresholds were deemed to exhibit a significant response to a given lever. We then examined how these significant responses were distributed across the 3 levers in each block using the binominal cumulative distribution function in Matlab. This calculated the probability that the number of neurons with an absolute t-stat value on a given lever above the 95^th^ percentile of the total distribution was greater than would be expected by chance, assuming that that significant t-stat values occurred at a rate of 10%, as noted above.

In order to track lever press responses on a trial-by-trial basis, it was necessary that there be a spike count value for all levers on all trials. The spike counts for each neuron were first normalized between 0 and 1. Interpolated vectors were then created so that there were both rewarded and nonrewarded normalized spike count values on every trial. PCA was performed separately on the matrix of normalized, interpolated spike count vectors for rewarded and nonrewarded (with ‘bias’ and ‘other’ levers combined) levers and the primary PCs were plotted. In addition, neurons were sorted based on the difference in normalized, interpolated spike counts in blocks 1 and 2 and the 100 neurons at the top and bottom of the distributions were submitted to k-means clustering. For k-means, a k of 3 was chosen because it allowed us to capture different dynamics in the most evenly enriched clusters (cluster sizes ranged from 27 to 41 neurons). Finally, a two-sample Kolmogorov-Smirnov tests was performed on each neuron to assess whether a significant change had occurred during the baseline periods (all timebins excluding those 1s prior to levers-out to 1s after the outcome) of blocks 1 versus 2 with a p-value corrected to 0.002.

## References

Addicott MA, Pearson JM, Sweitzer MM, Barack DL, Platt ML. (2017) A Primer on Foraging and the Explore/Exploit Trade-Off for Psychiatry Research. Neuropsychopharmacology. 42(10):1931–1939.

Aston-Jones G, Cohen JD. (2005) An integrative theory of locus coeruleus-norepinephrine function: adaptive gain and optimal performance. Annu Rev Neurosci. 28:403–50.

Azab H, Hayden BY. (2018) Correlates of economic decisions in the dorsal and subgenual anterior cingulate cortices. Eur J Neurosci. 47(8):979–993.

Azab H, Hayden BY. (2017) Correlates of decisional dynamics in the dorsal anterior cingulate cortex. PLoS Biol. 15(11):e2003091.

Barnett (2005) The Ethology of Domestic Animals: An Introductory Text. Ed. Jensen, P.

Basten U, Biele G, Heekeren HR, Fiebach CJ.(2010) How the brain integrates costs and benefits during decision making. Proc Natl Acad Sci U S A. 107(50):21767–72.

Bernacchia A, Seo H, Lee D, Wang XJ.(2011) A reservoir of time constants for memory traces in cortical neurons. Nat Neurosci. 14(3):366–72.

Blanchard TC, Hayden BY.(2014) Neurons in dorsal anterior cingulate cortex signal postdecisional variables in a foraging task. J Neurosci. 2014 Jan 8;34(2):646–55.

Bowden EM, Jung-Beeman M, Fleck J, Kounios J. (2005) New approaches to demystifying insight. Trends Cogn Sci. 9(7):322–8.

Cai X, Padoa-Schioppa C. (2012) Neuronal encoding of subjective value in dorsal and ventral anterior cingulate cortex. J Neurosci. 32(11):3791–808.

Caracheo BF, Emberly E, Hadizadeh S, Hyman JM, Seamans JK. (2013) Abrupt changes in the patterns and complexity of anterior cingulate cortex activity when food is introduced into an environment. Front Neurosci. May, 7:74.

Caracheo BF, Grewal JJS, Seamans JK. (2018) Persistent Valence Representations by Ensembles of Anterior Cingulate Cortex Neurons. Front Syst Neurosci. Oct., 12:51.

Cohen JD, McClure SM, Yu AJ. (2007) Should I stay or should I go? How the human brain manages the trade-off between exploitation and exploration. Philos Trans R Soc Lond B Biol Sci. 362(1481):933–42.

Cowen SL, Davis GA, Nitz DA. (2012) Anterior cingulate neurons in the rat map anticipated effort and reward to their associated action sequences. J Neurophysiol. 107(9):2393–407.

Cowen SL, McNaughton BL. (2007) Selective delay activity in the medial prefrontal cortex of the rat: contribution of sensorimotor information and contingency. J Neurophysiol. 98(1):303–16.

Critchley HD, Mathias CJ, Josephs O, O’Doherty J, Zanini S, Dewar BK, Cipolotti L, Shallice T, Dolan RJ. (2003) Human cingulate cortex and autonomic control: converging neuroimaging and clinical evidence. Brain. 126(Pt 10):2139–52.

Daw ND, O’Doherty JP, Dayan P, Seymour B, Dolan RJ. (2006) Cortical substrates for exploratory decisions in humans. Nature. 441(7095):876–9.

de Berker AO Kurth-Nelson Z, Rutledge RB, Bestmann S, Dolan RJ. (2019) Computing Value from Quality and Quantity in Human Decision-Making. J Neurosci. 39(1):163–176.

Durstewitz D, Vittoz NM, Floresco SB, Seamans JK. (2010) Abrupt transitions between prefrontal neural ensemble states accompany behavioral transitions during rule learning. Neuron. 66(3):438–48.

Euston DR, McNaughton BL. (2006) Apparent encoding of sequential context in rat medial prefrontal cortex is accounted for by behavioral variability. J Neurosci. 26(51):13143–55.

Feldman S, Conforti N. (1985) Modifications of adrenocortical responses following frontal cortex simulation in rats with hypothalamic deafferentations and medial forebrain bundle lesions. Neuroscience. 15(4):1045–7.

Groenewegen HJ, Uylings HB. (2000) The prefrontal cortex and the integration of sensory, limbic and autonomic information. Prog Brain Res. 126:3–28.

Groenewegen HJ, Wright CI, Uylings HB. (1997) The anatomical relationships of the prefrontal cortex with limbic structures and the basal ganglia. J Psychopharmacol. 11(2):99–106.

Hart Y, Mayo AE, Mayo R, Rozenkrantz L, Tendler A, Alon U, Noy L. (2017) Creative foraging: An experimental paradigm for studying exploration and discovery. PLoS One. 12(8):e0182133.

Hunt LT, Malalasekera WMN, de Berker AO, Miranda B, Farmer SF, Behrens TEJ, Kennerley SW. (2018) Triple dissociation of attention and decision computations across prefrontal cortex. Nat Neurosci. 21(10):1471–1481

Hayden BY, Pearson JM, Platt ML. (2011) Neuronal basis of sequential foraging decisions in a patchy environment. Nat Neurosci. 14(7):933–9.

Hayden BY, Platt ML. (2010) Neurons in anterior cingulate cortex multiplex information about reward and action. J Neurosci. 30(9):3339–46.

Hillman KL, Bilkey DK. (2010) Neurons in the rat anterior cingulate cortex dynamically encode cost-benefit in a spatial decision-making task. J Neurosci. 30(22):7705–13.

Hyman JM, Holroyd CB, Seamans JK. (2017) A Novel Neural Prediction Error Found in Anterior Cingulate Cortex Ensembles. Neuron. 95(2):447–456.

Hyman JM, Ma L, Balaguer-Ballester E, Durstewitz D, Seamans JK. (2012) Contextual encoding by ensembles of medial prefrontal cortex neurons. Proc Natl Acad Sci U S A. 109(13):5086–91.

Kennerley SW, Dahmubed AF, Lara AH, Wallis JD. (2009) Neurons in the frontal lobe encode the value of multiple decision variables. J Cogn Neurosci. 21(6):1162–78.

Khani A, Rainer G. (2016) Neural and neurochemical basis of reinforcement-guided decision making. J Neurophysiol. 116(2):724–41.

Klein-Flügge MC, Kennerley SW, Friston K, Bestmann S. (2016) Neural Signatures of Value Comparison in Human Cingulate Cortex during Decisions Requiring an Effort-Reward Trade-off. J Neurosci. 36(39):10002–1.

Kolling N, Akam T. (2017) (Reinforcement?) Learning to forage optimally. Curr Opin Neurobiol. 46:162–169.

Kolling N, Behrens T, Wittmann MK, Rushworth M. (2016) Multiple signals in anterior cingulate cortex Curr Opin Neurobiol. 37:36–43.

Kolling N, Behrens TE, Mars RB, Rushworth MF. (2012) Neural mechanisms of foraging. Science. 336(6077):95–8.

Lindsay AJ, Caracheo BF, Grewal JJS, Leibovitz D, Seamans JK. (2018) How Much Does Movement and Location Encoding Impact Prefrontal Cortex Activity? An Algorithmic Decoding Approach in Freely Moving Rats. eNeuro. Apr 27;5(2).

Ma L, Hyman JM, Durstewitz D, Phillips AG, Seamans JK. (2016) A Quantitative Analysis of Context-Dependent Remapping of Medial Frontal Cortex Neurons and Ensembles. J Neurosci. 36(31):8258–72.

Malagon-Vina H, Ciocchi S, Passecker J, Dorffner G, Klausberger T. (2018) Fluid network dynamics in the prefrontal cortex during multiple strategy switching. Nat Commun. 9(1):309.

Mobbs D, Hassabis D, Yu R, Chu C, Rushworth M, Boorman E, Dalgleish T. (2013) Foraging under competition: the neural basis of input-matching in humans. J Neurosci. 33(23):9866–72.

Monosov IE. (2017) Anterior cingulate is a source of valence-specific information about value and uncertainty. Nat Commun. 8(1):134.

Narayanan NS, Cavanagh JF, Frank MJ, Laubach M. (2013) Common medial frontal mechanisms of adaptive control in humans and rodents. Nat Neurosci. 16(12):1888–1895.

Neafsey EJ.(1990) Prefrontal cortical control of the autonomic nervous system: anatomical and physiological observations. Prog Brain Res. 85:147–65.

Pearson JM, Watson KK, Platt ML. (2014) Decision making: the neuroethological turn. Neuron. 82(5):950–65.

Powell NJ, Redish AD. (2016) Representational changes of latent strategies in rat medial prefrontal cortex precede changes in behaviour. Nat Commun. 7:12830.

Powell NJ, Redish AD.(2014) Complex neural codes in rat prelimbic cortex are stable across days on a spatial decision task. Front Behav Neurosci. Apr., 8:120.

Pratt WE, Mizumori SJ. (2001) Neurons in rat medial prefrontal cortex show anticipatory rate changes to predictable differential rewards in a spatial memory task. Behav Brain Res. 123(2):165–83.

Rushworth MF, Behrens TE. (2008) Choice, uncertainty and value in prefrontal and cingulate cortex. Nat Neurosci. 11(4):389–97.

Rushworth MF, Kolling N, Sallet J, Mars RB. (2012) Valuation and decision-making in frontal cortex: one or many serial or parallel systems? Curr Opin Neurobiol. 22(6):946–55.

Schultz W. (2015) Neuronal Reward and Decision Signals: From Theories to Data. Physiol Rev. 95(3):853–951.

Seo H, Lee D. (2007) Temporal filtering of reward signals in the dorsal anterior cingulate cortex during a mixed-strategy game. J Neurosci. 27(31):8366–77.

Sharpe MJ, Schoenbaum G.(2018) Evaluation of the hypothesis that phasic dopamine constitutes a cached-value signal. Neurobiol Learn Mem. 153(Pt B):131–136.

Stauffer WR, Lak A, Schultz W. (2014) Dopamine reward prediction error responses reflect marginal utility. Curr Biol. 24(21):2491–500.

Sutton RS & Barto AG (2018) Reinforcement Learning: An Introduction. Second Edition, MIT Press, Cambridge, MA.

Van Eden CG, Buijs RM. (2000) Functional neuroanatomy of the prefrontal cortex: autonomic interactions. Prog Brain Res. 126:49–62.

Wallis JD, Kennerley SW. (2011) Contrasting reward signals in the orbitofrontal cortex and anterior cingulate cortex. Ann N Y Acad Sci. 1239:33–42.

Wittmann MK, Kolling N, Akaishi R, Chau BK, Brown JW, Nelissen N, Rushworth MF. (2016) Predictive decision making driven by multiple time-linked reward representations in the anterior cingulate cortex. Nat Commun. 7:12327.

